# Highly Pathogenic Avian Influenza H5N5 in a Polar Bear and Atlantic Walrus, Svalbard, 2026, with Widespread Seroconversion in Polar Bears

**DOI:** 10.64898/2026.07.20.739480

**Authors:** Knut Madslien, Johanna Hol Fosse, Jon Aars, Cathrine Arnason Bøe, Magnus Andersen, Kayla Buhler, Ingvild Fjeldheim, Britt Gjerset, Torild Jørgensen, Ida Kristin Myhrvold, Andreas Rohringer, Kjersti Sturød, Morten Tryland, Bjørnar Ytrehus, Ragnhild Tønnessen, Ingebjørg Helena Nymo

## Abstract

Highly pathogenic avian influenza virus (HPAIV) subtype H5N5 was detected in a one-year-old polar bear (*Ursus maritimus*) and an adjacent adult Atlantic walrus (*Odobenus rosmarus rosmarus*), both found deceased in Raudfjorden, Svalbard. This represents the first confirmed case of HPAI in a European polar bear and the second in an Atlantic walrus. Viral genomes were nearly identical and harbored PB2-E627V, a marker associated with mammalian adaptation. Several polar bears, including the deceased individual, had previously been observed feeding on the walrus carcass. Antibodies against H5 were detected in 75% of polar bears in 2023 (n=36) and 97% in 2024-2025 (n=65), suggesting extensive circulation of HPAIV in the population following the first detections in birds in Svalbard in 2022, whereas no antibodies were detected in samples from 2014-2022 (n=243).

## Introduction

Highly pathogenic avian influenza (HPAI) has emerged as a major global health threat. Since 2020, HPAI viruses (HPAIVs) of the H5 clade 2.3.4.4b have driven an ongoing panzootic with widespread circulation in wild birds across multiple continents (1), including recent spread to Antarctica (2) and Australia (3). Increasing spillover to mammals (4) raises concerns about viral adaptation and cross-species transmission.

HPAIV was first detected in the high Arctic Archipelago of Svalbard in 2022 in a glaucous gull (*Larus hyperboreus*) and has since been detected in multiple bird species (5), Atlantic walrus (*Odobenus rosmarus rosmarus*) (6) and Arctic foxes (*Vulpes lagopus*) (5). Although Arctic ecosystems are geographically remote, they are tightly connected to ecosystems across continents through migratory birds (5). Arctic wildlife may be particularly vulnerable due to low pre-existing immunity (7), strong seasonal aggregation (8), and environmental conditions that favor virus persistence (9), but limited surveillance hampers understanding of virus dynamics.

Polar bears (*Ursus maritimus*) in Svalbard belong to the Barents Sea subpopulation shared by Norway and the western Russian Arctic. The population recovered after hunting was banned in 1973 and is currently stable or increasing, despite distributional shifts associated with sea-ice loss (10). The bears are generally in good body condition (11).

Fatal infection with HPAIV H5N1 has been described in a free-ranging juvenile polar bear at Point Barrow, Alaska, USA, in 2023 (12). In addition, HPAIV H5N5 has been detected in a polar bear in 2025 based on publicly available sequence data (GISAID: EPI_ISL_20423927) (13). Antibodies to influenza A virus (IAV) were not detected in polar bears (n=100) from the Arctic Coastal Plain in Alaska during 2013-2016 (7). In a survey of 25 polar bears in Franz Josef Land and Novaya Zemlya, Russia, in 2010-2011, two adults were seropositive for IAV, whereas nine juveniles and cubs were seronegative (14). Natural HPAIV H5N1 infections have also been documented in other bear species (12).

Svalbard Atlantic walruses are part of a shared Svalbard–Franz Josef Land population, nearly driven to extinction by past harvesting but now recovering following protection in 1952 (15, 16).

HPAIV H5N5 was detected in a Svalbard Atlantic walrus in 2023 (6). Anti-IAV antibodies were not detected in Atlantic walruses (n=210) from the eastern high Canadian Arctic sampled in 1984-1998 (17), but antibodies were detected in eight Pacific walruses (*Odobenus rosmarus divergens*, n=38) from St. Lawrence Island and Round Island, Alaska, sampled in 1994-1996 (18).

We report detection of HPAIV H5N5 in a polar bear and an Atlantic walrus in Svalbard, Norway, in May 2026, together with evidence of population-level seroconversion to H5 avian influenza virus in the polar bear population in 2023, followed by high seroprevalence.

## Methods

### Observations of Sick Polar Bears and Carcasses

Field observations of sick and dead animals were reported to the Norwegian Polar Institute (NPI) and the Norwegian Veterinary Institute (NVI) by local tourist guides. Information on carcass location, timing, and animal behavior was compiled to guide field investigations and sampling.

### Sample Collection from Polar Bear and Walrus Carcasses

In response to suspected notifiable disease (i.e. HPAI or rabies) (19), NVI wildlife veterinarians assisted the Governor of Svalbard with sample collection on May 14. The carcasses were located approximately 150 m apart (Figure 1), the bear in lateral (Figure 2) and the walrus in sternal recumbency (Figure 3). Polar bear footprints and bird feces in the snow indicated high scavenger activity. The condition of the carcasses, challenging weather, and personnel safety risks posed by nearby polar bears precluded a complete field necropsy. Tissue samples from parotid salivary glands, liver, and brain, and swabs from the nostrils, brain, trachea, and parotid salivary glands were collected from the polar bear and stored in virus transport medium (VTM, Copan Universal Transport Medium, Copan Diagnostics, Murrieta, CA, USA) (20). Tissue samples from the brain and brain swabs were collected from the walrus and stored in VTM.

**Figure 1.**
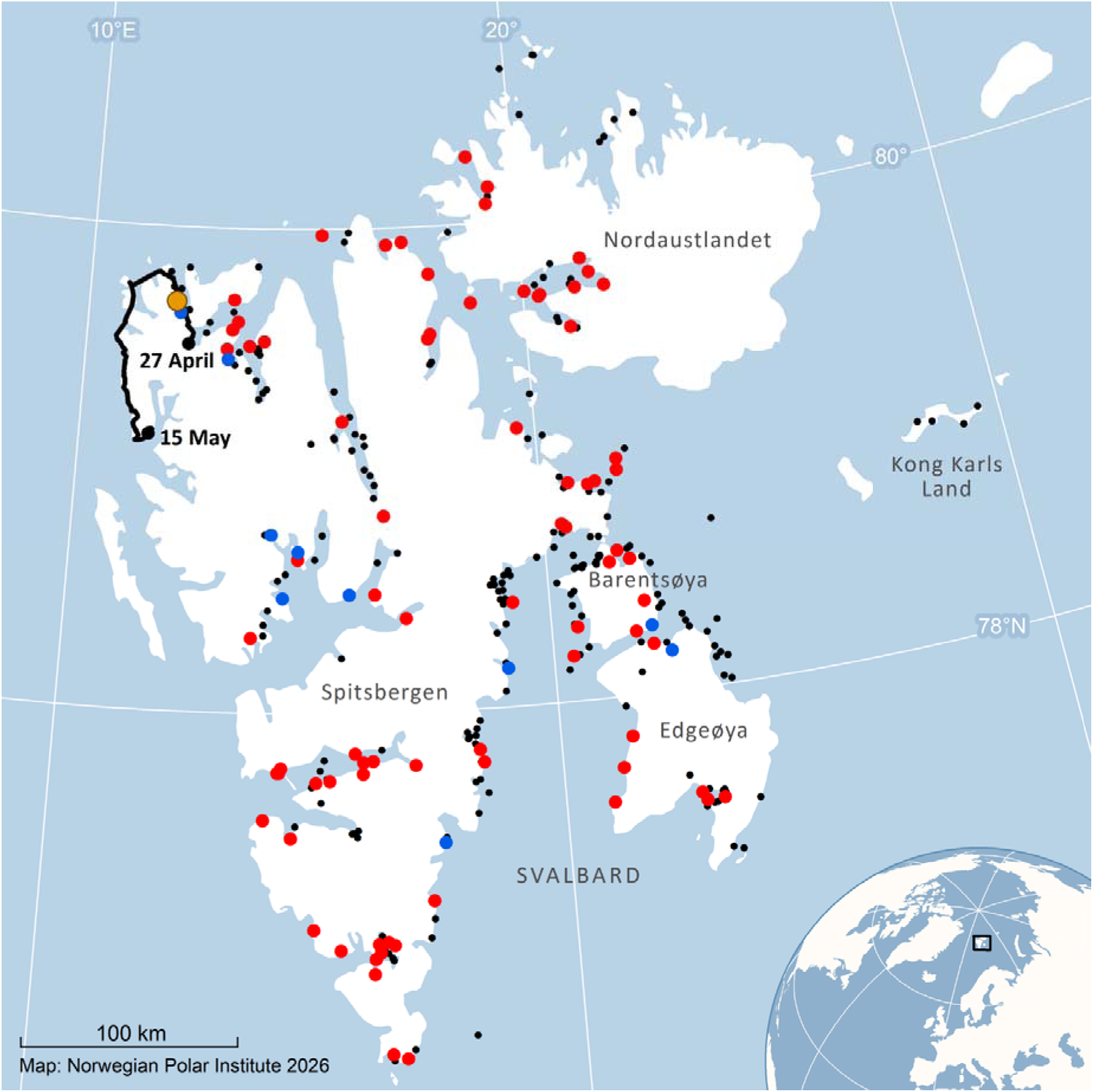
Map of Svalbard showing the locations of the HPAIV H5N5-positive walrus and polar bear carcasses in May 2026 (orange dots). The black line shows the movement track of the polar bear’s mother from April 27 to May 15, 2026. Black dots indicate capture locations of bears from 2014-2017 and 2022 (all H5 seronegative). For 2023-2025, blue dots indicate seronegative and red dots seropositive bears.

**Figure 2.**
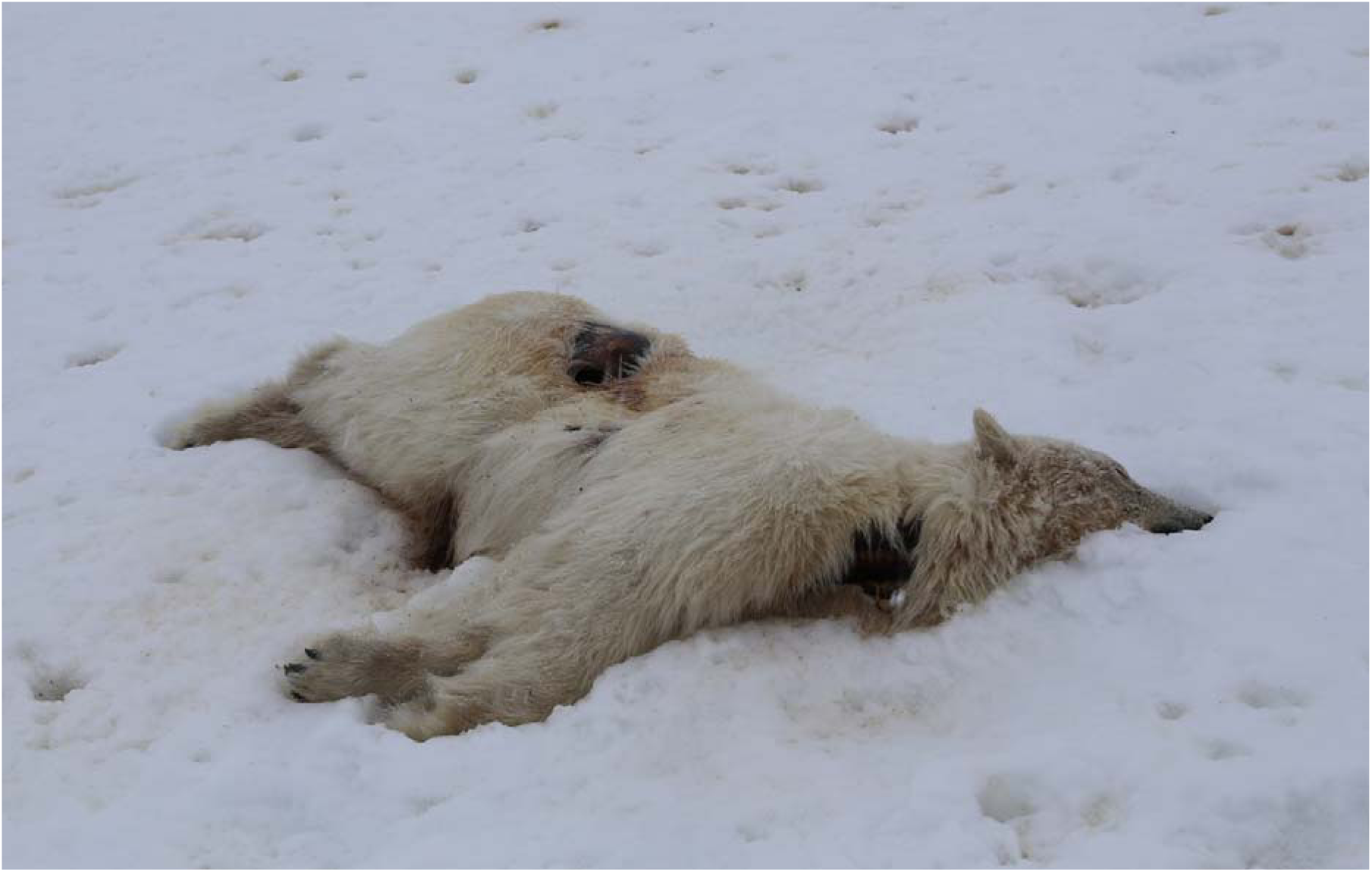
Highly pathogenic avian influenza virus (HPAIV) H5N5 was detected in a brain sample from the carcass of a yearling, male polar bear in Svalbard, Norway, in May 2026.

**Figure 3.**
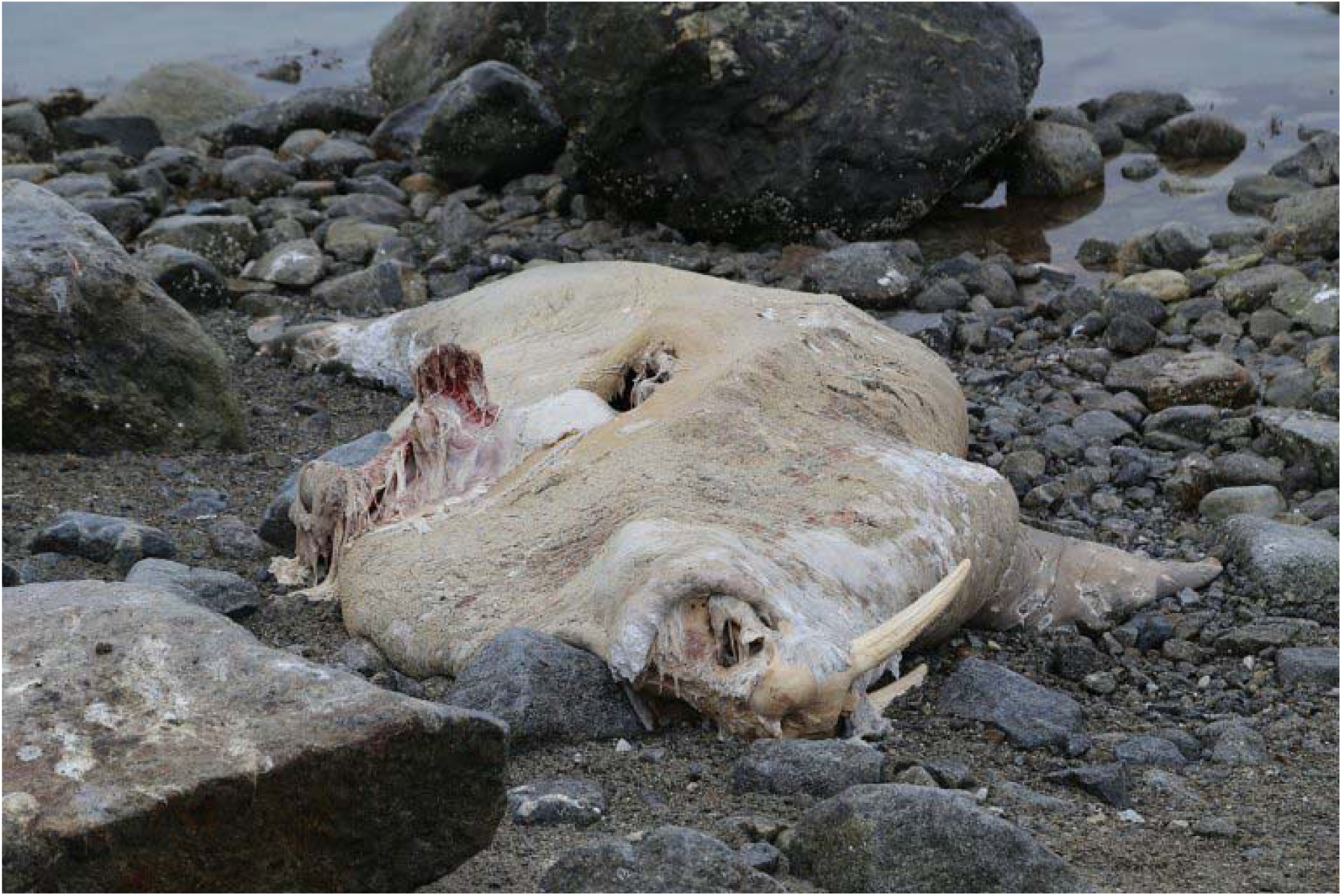
Highly pathogenic avian influenza virus (HPAIV) H5N5 was detected in a brain sample from the carcass of an adult, male Atlantic walrus in Svalbard, Norway, in May 2026.

### Virus Detection and Subtyping by real-time RT-PCR

Swab and tissue samples were pretreated with MagNA Pure External Lysis Buffer (Roche, Basel, Switzerland), and nucleic acids were extracted using the MagNA Pure 24 automated system (Roche) according to the manufacturer’s instructions. Samples were analyzed by real-time RT-PCR (rRT-PCR) targeting the nucleoprotein (N) gene of rabies virus (RABV) (21), and the M1 gene of IAV (22). IAV-positive samples were further characterized by rRT-PCR for hemagglutinin (HA) and neuraminidase (NA) subtyping and by an rRT-PCR assay targeting HPAIV H5 clade 2.3.4.4b (23, 24). IAV assays were performed using the OneStep RT-PCR Kit (Qiagen, Hilden, Germany), whereas RABV assays were conducted using the SuperScript III Platinum One-Step qRT-PCR Kit (Invitrogen, Thermo Fisher Scientific, Waltham, MA, USA). All analyses were carried out on AriaMX instruments (Agilent Technologies, Santa Clara, CA, USA).

### Phylogeny and Mutation Analysis

Whole genome sequencing (WGS) was performed as previously described (25). Consensus sequences were assembled using IRMA v1.2.0 (26). Sequences generated in this study are available in GISAID (13) under accession numbers EPI_ISL_20456296 (A/Polar_bear/Norway/2026-80-38-1-4/2026) and EPI_ISL_20456295 (A/Walrus/Norway/2026-80-37-1-4/2026). Sequences were genotyped using the GenIn2 and GenoFlu tools (GISAID platform). BLAST searches of the HA, NA, and PB2 gene segments of the polar bear virus (EPI_ISL_20456296) were conducted in GISAID EpiFlu. For each segment, the 100 closest matches were identified, and the corresponding genomes were retrieved. Previously generated sequences from Svalbard and the first (A/White-tailed_eagle/Norway/2022-07-100/2022) and most recent (A/Glaucous_gull/Norway/2025-07-2990-5-1/2025) Norwegian detections of EA-2021-I, were included. The reference sequence for the EA-2021-I genotype (EPI_ISL_1665268 A/whooper_swan/Romania/10123_21VIR849-1/2021) was used for rooting. Coding regions were aligned with MUSCLE in MEGA X v10.1.8 (27), curated, and trimmed to open reading frames. Partial and duplicate sequences were removed. Segments were concatenated (PB2/PB1/PA/HA/NP/NA/MP/NS). Phylogenies (PB2/HA/NA/concatenated) were inferred using maximum likelihood (Tamura-Nei model) (28) with 1000 bootstrap replicates. Ambiguous sites were excluded. All sequences included in the analyses and associated metadata are available at https://doi.org/10.55876/gis8.260612kn, whilst Appendix Table 1 contains sequence identifiers and metadata.

**Table 1.**
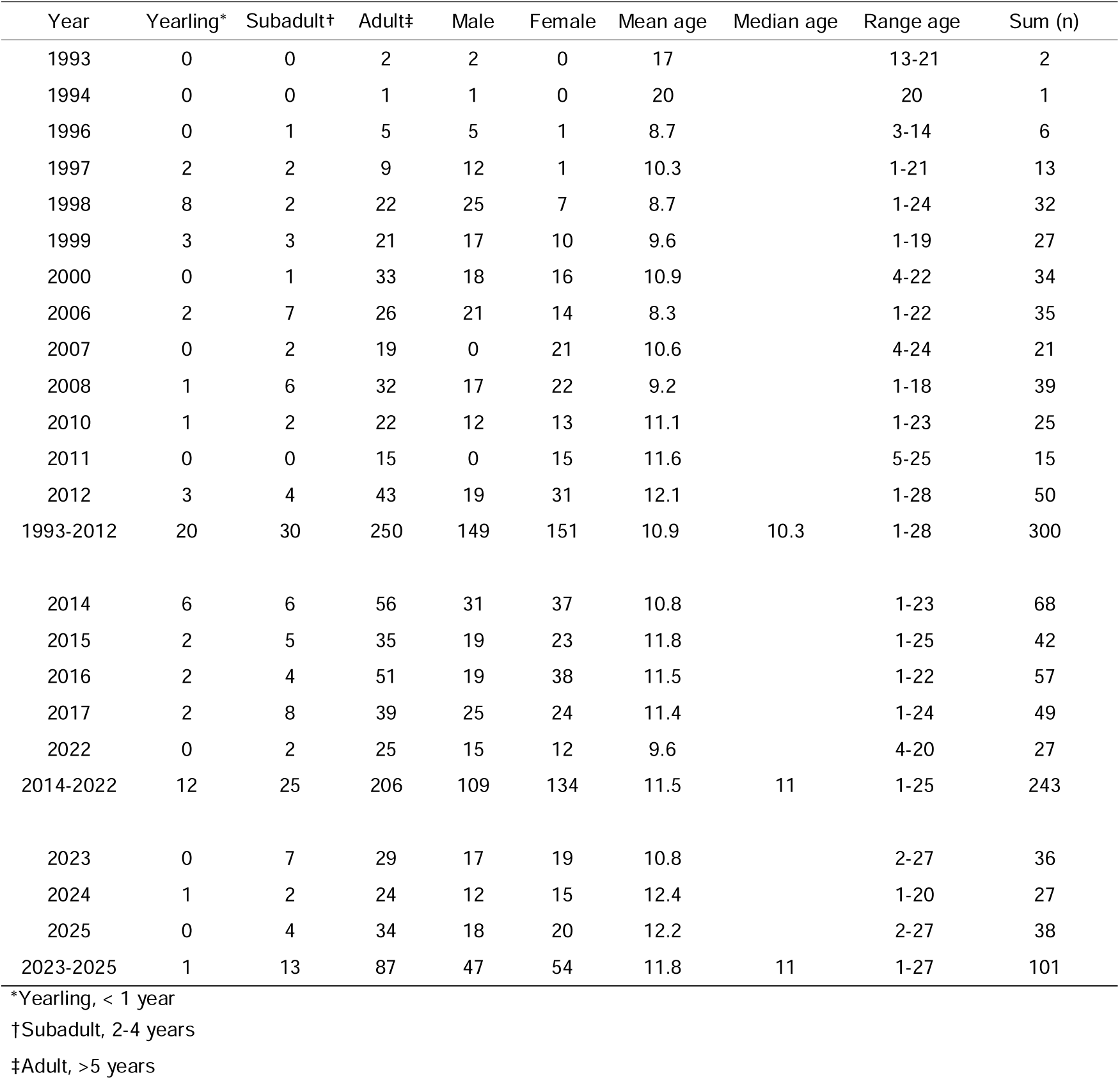
Blood samples from polar bears (n= 644), distributed by age-class, sex and age, collected in Svalbard between 1993 and 2025.

| Year | Yearling* | Subadult† | Adult‡ | Male | Female | Mean age | Median age | Range age | Sum (n) |
| --- | --- | --- | --- | --- | --- | --- | --- | --- | --- |
| 1993 | 0 | 0 | 2 | 2 | 0 | 17 |  | 13-21 | 2 |
| 1994 | 0 | 0 | 1 | 1 | 0 | 20 |  | 20 | 1 |
| 1996 | 0 | 1 | 5 | 5 | 1 | 8.7 |  | 3-14 | 6 |
| 1997 | 2 | 2 | 9 | 12 | 1 | 10.3 |  | 1-21 | 13 |
| 1998 | 8 | 2 | 22 | 25 | 7 | 8.7 |  | 1-24 | 32 |
| 1999 | 3 | 3 | 21 | 17 | 10 | 9.6 |  | 1-19 | 27 |
| 2000 | 0 | 1 | 33 | 18 | 16 | 10.9 |  | 4-22 | 34 |
| 2006 | 2 | 7 | 26 | 21 | 14 | 8.3 |  | 1-22 | 35 |
| 2007 | 0 | 2 | 19 | 0 | 21 | 10.6 |  | 4-24 | 21 |
| 2008 | 1 | 6 | 32 | 17 | 22 | 9.2 |  | 1-18 | 39 |
| 2010 | 1 | 2 | 22 | 12 | 13 | 11.1 |  | 1-23 | 25 |
| 2011 | 0 | 0 | 15 | 0 | 15 | 11.6 |  | 5-25 | 15 |
| 2012 | 3 | 4 | 43 | 19 | 31 | 12.1 |  | 1-28 | 50 |
| 1993-2012 | 20 | 30 | 250 | 149 | 151 | 10.9 | 10.3 | 1-28 | 300 |
| 2014 | 6 | 6 | 56 | 31 | 37 | 10.8 |  | 1-23 | 68 |
| 2015 | 2 | 5 | 35 | 19 | 23 | 11.8 |  | 1-25 | 42 |
| 2016 | 2 | 4 | 51 | 19 | 38 | 11.5 |  | 1-22 | 57 |
| 2017 | 2 | 8 | 39 | 25 | 24 | 11.4 |  | 1-24 | 49 |
| 2022 | 0 | 2 | 25 | 15 | 12 | 9.6 |  | 4-20 | 27 |
| 2014-2022 | 12 | 25 | 206 | 109 | 134 | 11.5 | 11 | 1-25 | 243 |
| 2023 | 0 | 7 | 29 | 17 | 19 | 10.8 |  | 2-27 | 36 |
| 2024 | 1 | 2 | 24 | 12 | 15 | 12.4 |  | 1-20 | 27 |
| 2025 | 0 | 4 | 34 | 18 | 20 | 12.2 |  | 2-27 | 38 |
| 2023-2025 | 1 | 13 | 87 | 47 | 54 | 11.8 | 11 | 1-27 | 101 |

\*Yearling, < 1 year
†Subadult, 2-4 years
‡Adult, >5 years

FluMutGUI v3.2.0 was used to screen the polar bear and walrus viruses for markers of mammalian adaptation (29). Sequences were compared with HPAIV H5N5 from birds and mammals in Svalbard and Jan Mayen (2022-2026; Appendix Table 2). Additional mutations were identified by comparison with Norwegian H5N5 sequences from 2022: A/White-tailed_eagle/Norway/2022-07-446-1/2022 and A/Northern_gannet/Norway/2022-07-1233-1/2022.

### Sampling and GPS-collaring of Live Polar Bears

On April 26, 2025, the deceased yearling (N26474) and its mother (N26240) were captured by NPI as part of the annual polar bear population monitoring program. Blood samples from N26240 and other Svalbard polar bears (n=644, Table 1) were collected during 1993-2025 (March 23-May 3). Bears were captured from a helicopter using tiletamine-zolazepam (5-10 mg/kg) or tiletamine-zolazepam-medetomidine (approx. 2.2 mg/kg and 0.06 mg/kg) following standard procedures (30). Age was estimated for adults from a vestigial premolar tooth extracted at first capture (31). Blood was collected from *V. femoralis*, centrifuged (3000 g, 10 min, room temperature), and serum or plasma was retrieved and frozen (-18 °C). Adult females were equipped with GPS collars (Argos satellite telemetry, CLS, Toulouse, France, via the Iridium network, 2 h sending frequency). The sampling was approved by the Norwegian Animal Research Authority (FOTS ID 31180).

### Antibody detection

Heat-inactivated plasma were analyzed for antibodies to H5 avian influenza (anti-H5, n=344) and IAV nucleoprotein (anti-NP, n=496) using commercial ELISAs (ID Screen® Influenza H5 Antibody Competition 3.0 Multi-species ELISA and ID Screen® Influenza A Antibody Competition Multi-species ELISA, Innovative Diagnostics, France), at dilutions of 1:2 and 1:10 respectively, following the manufacturer’s recommendations for ferrets. Anti-H5-positive samples were pretreated with receptor-destroying enzyme (3:1, 37 °C, overnight), cross adsorbed against chicken erythrocytes, and tested for hemagglutination inhibition (HI) using four hemagglutinating units of H5 clade 2.3.4.4b virus antigen (A/turkey/Italy/VIR9520-320/2021(H5N1)), with titers ≥ 1:20 interpreted as positive. Serology data are provided in Appendix Tables 3-5.

## Results

### Field Observations and Mortality Timeline

An adult walrus was first observed dead in Raudfjorden, Svalbard (79°41′N, 11°60′E; Figure 1), around April 11, 2026, and was considered relatively fresh. On April 28, a female polar bear (N26240) and her yearling cub (N26474) were observed feeding on the carcass. The female was identified visually by her GPS collar, and telemetry data confirmed that she arrived in the area on April 28. During the same period, two additional polar bears in the area were reported to show signs of lameness. The yearling cub was found dead on May 10, with the time of death estimated to May 8, after which the mother left the area. Telemetry data from the GPS-collared female (N26240) showed that the bears primarily occupied coastal areas of Prins Karls Forland and northwest Spitsbergen before the event (Figure 1).

### Gross Findings

The polar bear was male, approximately 17 months old, and in fair body condition with ear tags (N26474) from NPI. Field examination revealed no signs of intravital external trauma, but multiple openings into the body cavity. Several internal organs were missing, and the liver, lungs, and heart showed marked autolysis. The walrus was an adult male. Both carcasses were in an advanced state of decomposition, precluding full necropsy.

### Virus Detection and Characterization

HPAIV H5N5 clade 2.3.4.4b was detected by rRT-PCR in brain samples from both animals with low Cq values (Cq 18-19) indicating high virus loads, and in a nasal swab from the polar bear (Cq 33), indicating a lower viral load. Both animals tested negative for RABV. Complete HPAIV genomes were obtained by WGS. The polar bear and walrus viruses were nearly identical, differing at only three nucleotide positions across the 13,079-nt coding genome. Genotyping assigned both viruses to EA-2021-I (GenIn2) and A6 (GenoFlu), representing equivalent genotype classifications in the European and North American nomenclature frameworks. The three differing sites showed ∼20% intra-sample variation in the sequencing data and were therefore assigned ambiguity codes. At PB1 position 208, the major variant in the polar bear-derived sequence matched the walrus consensus sequence. The remaining differences occurred at two adjacent positions in the PA segment (1464–1465), where the walrus-derived sequence contained GT, and the polar bear-derived sequence contained AA. This difference resulted in an amino acid substitution at PA position 489 (C489S). Notably, the walrus-associated GT variant was also present as a minor variant (∼20%) in the polar bear virus population (Appendix Table 6).

Phylogenetic analyses (Figure 4; Appendix Figure 1) demonstrated that the viruses belonged to a well-supported lineage comprising detections in wild birds and mammals from Norway, Iceland, and Canada during 2024-2026. A defining feature of this lineage was PB2-E627V, a mutation associated with mammalian adaptation. This lineage was phylogenetically distinct from the branch containing the previously reported H5N5 virus from an Alaskan polar bear (EPI_ISL_20423927) and the H5N5 human case in Washington State, USA (32).

**Figure 4.**
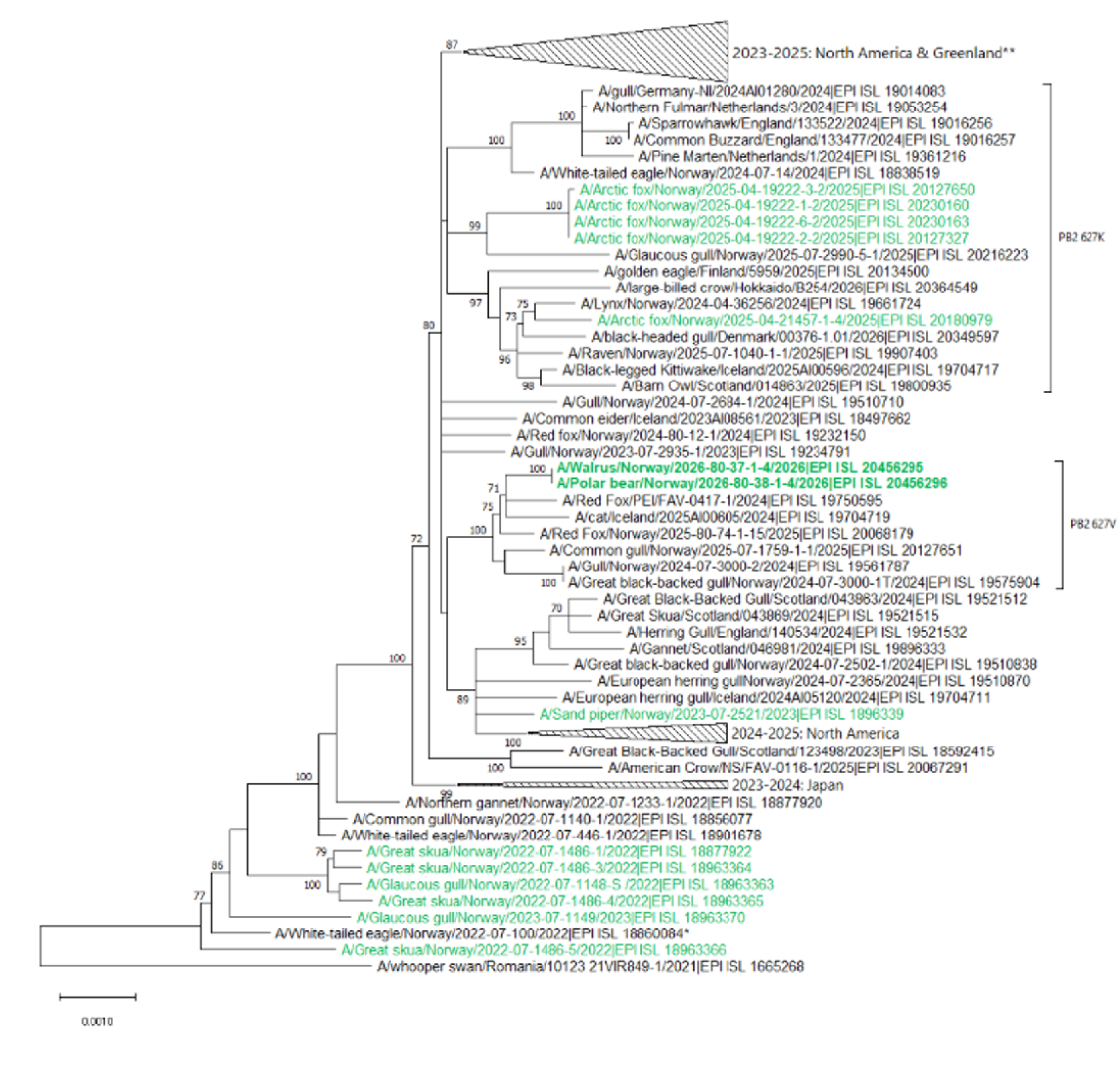
Phylogenetic characterization of the highly pathogenic avian influenza virus (HPAIV) H5N5 genomes obtained from a polar bear and a walrus carcass, Svalbard, May 2026. Rooted maximum-likelihood tree (Tamura-Nei) based on concatenated coding regions from a curated set of 97 sequences (all genotype EA-2021-I/A6). Bootstrap support values ≥70 are shown at nodes. Sequences from the walrus and polar bear are shown in bold, sequences from Svalbard are highlighted in green. Clades defined by amino acid substitutions at PB2 position 627 are indicated in brackets. Selected branches are collapsed for clarity. The complete tree, as well as separate trees for PB2, HA, and NA, are available in Appendix Figure 1 A-D. *First detection of genotype EA-2021-I (H5N5) in Norway. **Collapsed branch includes the first reported H5N5 detections in a human and a polar bear.

FluMut analysis also identified several other substitutions previously reported as markers of mammalian adaptation. However, most were present in both mammalian and avian viruses from the lineage (Appendix Table 2) and are commonly observed in contemporary H5 clade 2.3.4.4b viruses. No substitutions associated with altered receptor-binding specificity were detected in HA, indicating retention of a predominantly avian-like receptor-binding profile. Likewise, no lineage-specific M1 or NS1 substitutions uniquely associated with mammalian viruses were identified.

Comparative sequence analysis identified several lineage-associated substitutions, including PB1-I181M and NP-V105M, which co-occurred with PB2-E627V. Additional substitutions, HA-A172T and NS1-R59H, were detected in recent H5N5 viruses collected during 2025–2026. In contrast, PB2-V255F and NS-L33R were detected only in sequences from the polar bear and walrus. The N5 neuraminidase contained a 22-aa stalk deletion at positions 47–68, a characteristic feature of this H5N5 lineage that has been present since 2021 (33), and previously associated with altered glycosylation, increased virulence, and host adaptation (34). No markers of antiviral resistance were identified.

### Antibody detection

Anti-H5 antibodies were not detected in polar bears sampled in 2014-2017 and 2022 but were detected in 27/36 polar bears sampled in 2023 (75%, 95% CI 58-88%), 26/27 in 2024 (96%, 95% CI 81-100%), and 37/38 in 2025 (97%, 95% CI 86-100%) (Figure 5A). Longitudinal sampling showed seroconversion within individuals. Bear N26240 seroconverted between 2023 and 2024 (Figure 5B). Among anti-H5 positive samples, HI titers of ≥1:20 were detected in 7/27 samples in 2023 (1:20-1:40), 11/26 in 2024 (1:20-1:320), and 16/37 in 2025 (1:20-1:40) (Figure 5C-D). HI titers in individual bears appeared to be transient, and N26240 was HI negative (Appendix Table 5). Anti-NP followed a similar pattern (Figure 5E), with antibodies detected in 21/36 samples in 2023, 22/27 in 2024, and 31/38 in 2025, compared with 1/68 in 2014 and 0/27 in 2022. No anti-NP was detected in a complementary historical cohort (1993-2012, n=300).

**Figure 5A.**
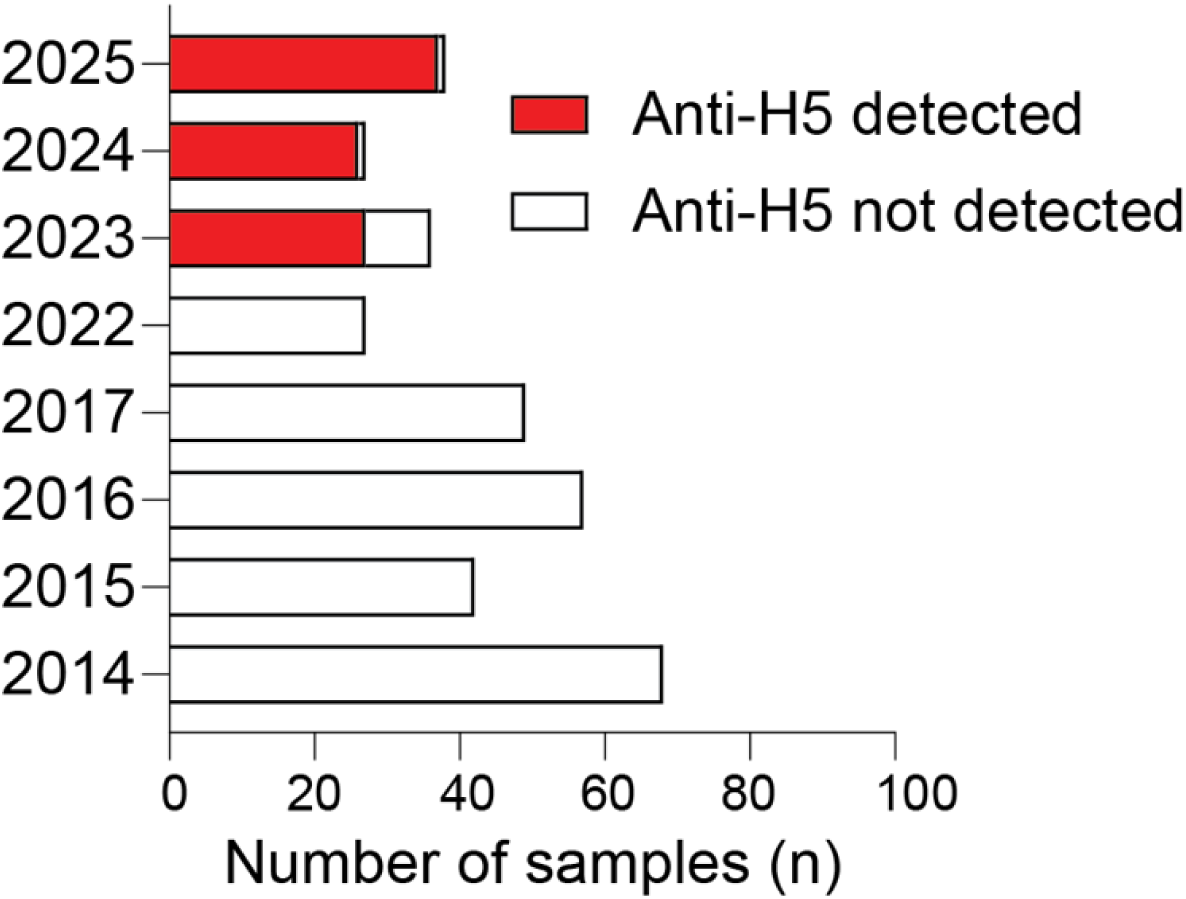
Serology reveals a population-level anti-H5 seroconversion in Svalbard polar bears between April 2022 and March 2023. Number of anti-H5 positive (red) and negative (white) samples by ELISA, collected from polar bears in Svalbard in 2014-2017 and 2022-2025.

**Figure 5B.**
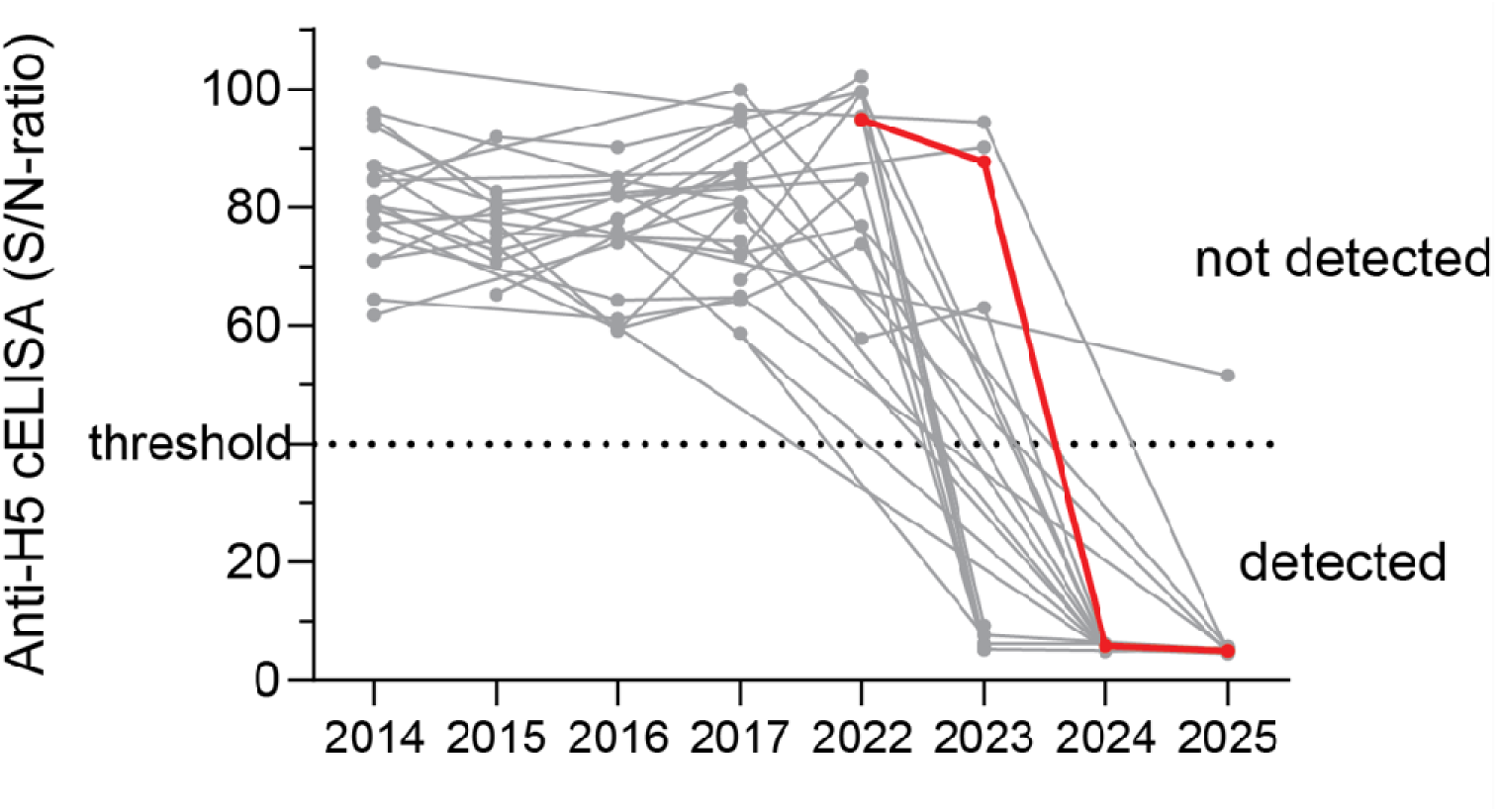
Serology reveals a population-level anti-H5 seroconversion in Svalbard polar bears between April 2022 and March 2023. Longitudinal sampling between 2014 and 2025 illustrates anti-H5 seroconversion in individuals, showing those with ≥three samples. Dots mark S/N-values of each sample, and lines connect values from the same individual. Red denotes mother of deceased yearling. S/N<40 = positive.

**Figure 5C.**
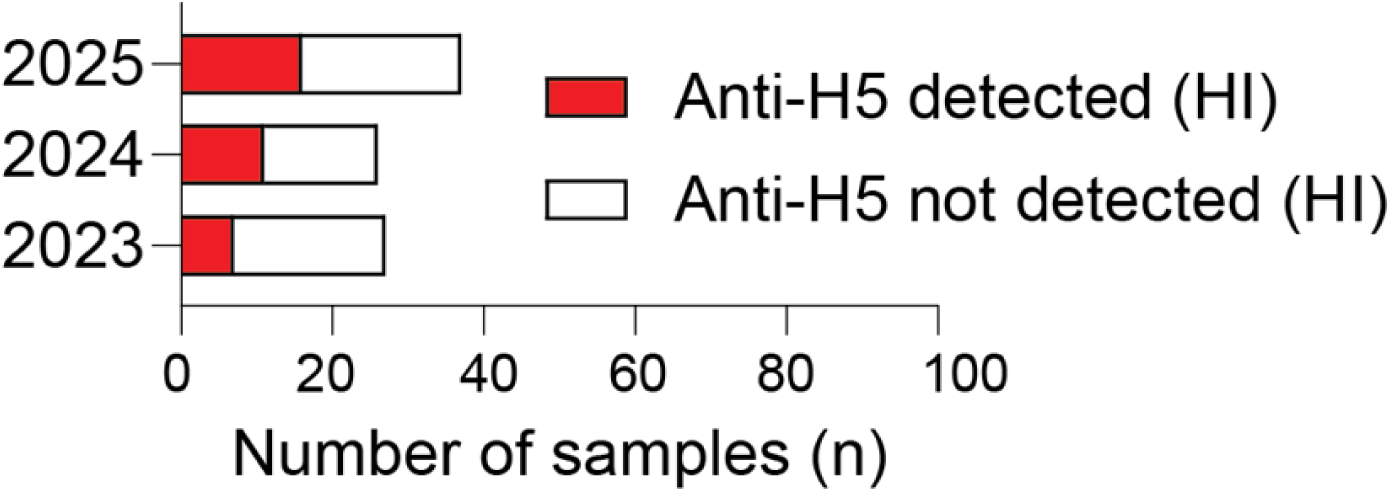
Serology reveals a population-level anti-H5 seroconversion in Svalbard polar bears between April 2022 and March 2023. Number of anti-H5 positive samples (Figure 5A) testing positive (red) and negative (white) in HI test with a H5 clade 2.3.4.4b antigen, collected from polar bears from Svalbard between 2023 and 2025.

**Figure 5D.**
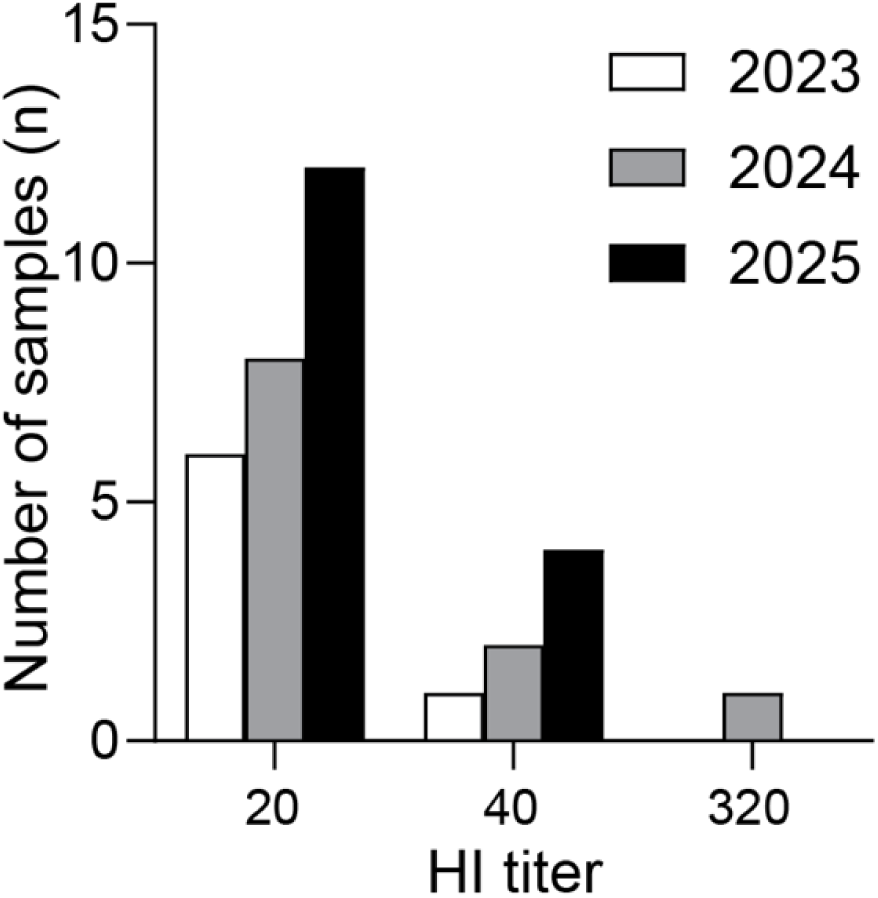
Serology reveals a population-level anti-H5 seroconversion in Svalbard polar bears between April 2022 and March 2023. Distribution of titers among HI positive samples, collected from polar bears in Svalbard between 2023 and 2025.

**Figure 5E.**
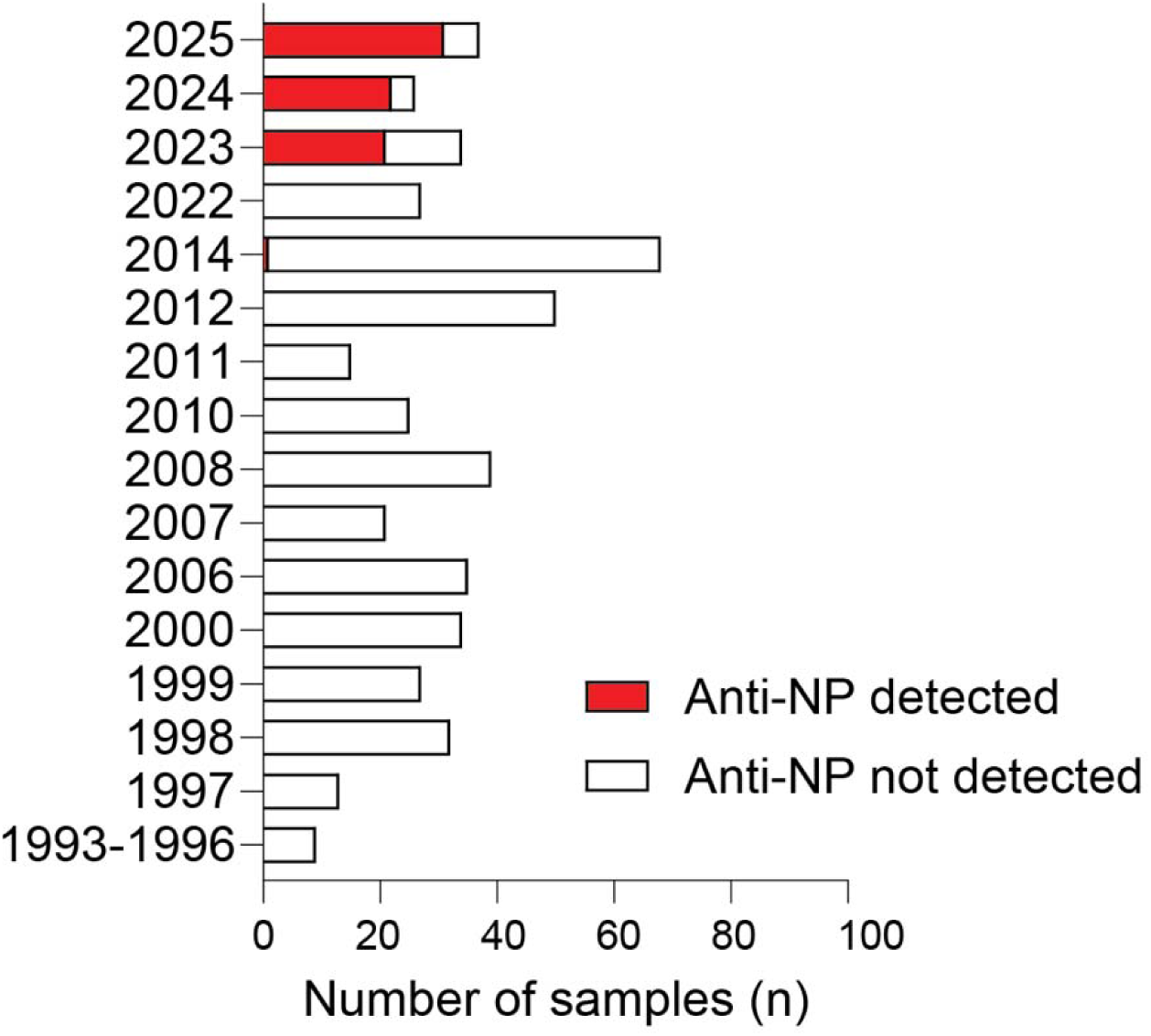
Serology reveals a population-level anti-H5 seroconversion in Svalbard polar bears between April 2022 and March 2023. Samples (n=644) were collected from polar bears in Svalbard in 1993-2012 (n = 300), 2014-2022 (n = 243), 2023 (n = 36), 2024 (n = 27), and 2025 (n = 38). Number of anti-influenza A nucleoprotein (anti-NP) positive (red) and negative (white) by ELISA. Complete data provided in Appendix Tables 3-5.

## Discussion

We report the first confirmed case of HPAIV in a polar bear in Europe and the second (6) in an Atlantic walrus. The near-identical viral genomes, including the mammalian-associated PB2-E627V substitution, and close spatiotemporal overlap indicate an epidemiological link. This is supported by observations of the deceased polar bear scavenging the walrus carcass and telemetry data showing that it was present at the site over ten days.

A high proportion of Svalbard polar bears seroconverted to anti-H5 after the sampling season in spring 2022, coinciding in time with the first detection of HPAIV H5Nx clade 2.3.4.4b in local seabirds during summer 2022 (5). The first seropositive polar bears were detected in 2023, with seroprevalence >95% in 2024–2025. Longitudinal sampling confirmed seroconversion in individuals during this period. Only a proportion yielded positive HI titers, mostly in the low range and transient. Together, these findings provide evidence for a recent transition from an immunologically naïve population to one where most individuals have been exposed to H5 IAV.

The high seroprevalence suggests that many exposed polar bears survive infection with current H5 HPAIV strains. However, clinical disease can occur, as demonstrated by the present case, the previous confirmed case (12), and detection of virus in brain tissue. Observations of lameness in other bears in the same area as the deceased yearling support this, although infection was not confirmed. The mother of the infected yearling was seropositive since 2024 but tested negative by HI. She left the area on May 12 in apparently good physical condition despite likely exposure, possibly reflecting partial protection associated with previous infection. The Barents Sea polar bear subpopulation increased after hunting was banned in 1973 and is now stable or increasing (10, 11), suggesting that HPAI has not, to date, affected population size.

The walrus carcass was relatively fresh when first observed, and bears may have been infected by scavenging the carcass. Under Arctic conditions, low temperatures may prolong viral persistence in carcasses (35). For the walrus, the source of infection remains uncertain but may involve exposure to contaminated environments, such as mussels where IAV has been detected earlier (36), or contact with infected seabirds (37, 38). The two detections of HPAIV H5N5 in walruses on Svalbard (6), without concurrent observations in local bird populations, raise the possibility that marine mammals may play an overlooked role in the epidemiology of this subtype.

The detection of PB2-E627V in the polar bear and walrus viruses is notable. Among the HP H5N5 genomes currently available in GISAID (n = 517 as of May 31, 2026), this mutation is exclusively found within the phylogenetic lineage that includes polar bear and walrus viruses (n = 9). Other members of this branch include viruses detected in red foxes (Canada 2024, Northern Norway 2025), a cat (Iceland 2025), and avian hosts, particularly gulls (mainland Norway 2024-2025, Iceland 2025). Its occurrence across multiple species and geographically distinct regions over time suggests maintenance within a circulating H5N5 lineage, rather than repeated independent emergences.

A similar pattern of PB2-E627K in a distinct phylogenetic branch indicates diversification at this key host-range determinant. PB2 is a subunit of the IAV RNA polymerase complex (39), and substitutions at position 627 influence host adaptation, with E627K marking enhanced replication in mammalian cells (40). PB2-E627V may confer a more intermediate phenotype allowing efficient viral replication in both avian and mammalian hosts (41). The detection of this substitution in multiple mammalian species, including marine mammals, suggests recurring opportunities for infection within Arctic ecosystems. This substitution alone is unlikely to fully account for efficient host adaptation and likely acts in combination with additional, poorly characterized mutations in recent H5N5 viruses to influence viral fitness. In addition to PB2-E627V, consistent substitutions were identified in other components of the viral polymerase complex, including PB1-I181M and NP-V105M, which co-occur in recent viruses and appear to be established within this lineage. This pattern suggests coordinated changes affecting polymerase function rather than reliance on a single host-adaptation marker.

No substitutions associated with altered receptor-binding specificity were identified in HA, indicating retention of an avian-like receptor-binding profile. Similarly, no lineage-specific changes in M1 or NS1 linked to mammalian adaptation were observed. Sub-consensus variation at PA position 489 may indicate intra-host diversification within the polymerase complex. This site has been implicated in brain tissue in experimental H5N1 infections (42), suggesting potential biological relevance.

Although no human cases have been linked to the HPAIV H5N5 PB2-E627V cluster, its detection across diverse mammalian species indicates a broad host range. Occurrence in both terrestrial and marine mammals suggests that ecological interactions, including predation and scavenging, may facilitate cross-species transmission. These findings are consistent with incremental, multigenic adaptation enabling infection of multiple hosts without evidence of full mammalian adaptation. Although HPAIV H5N5 can be zoonotic (43), the risk of zoonotic spillover to humans is considered very low for H5N5 detected in wildlife species with little or no human contact. Nevertheless, the transmission to novel host species underscores the importance of timely risk assessment, surveillance, and infection preparedness, as well as awareness and adherence to appropriate biosafety measures among wildlife professionals. Continued circulation within an avian-associated lineage as well as uncertainties on disease reservoir and adaptation underscores the need for disease surveillance at the wildlife–human interface in Arctic and North Atlantic regions.

The current report highlights that H5 HPAIV circulates in Arctic ecosystems and infects apex predators. Although no population-level effects are currently evident, detection in brain tissue and field observations suggest that disease can occur at the individual level. The identification of PB2-E627V and additional polymerase-associated changes is consistent with adaptation to mammals within an avian-associated lineage. These findings highlight the need for continued surveillance to clarify transmission pathways, evolutionary dynamics, and potential health effects.

## Acknowledgements

We gratefully acknowledge all data contributors, i.e., the authors and their originating laboratories responsible for obtaining the specimens, and their submitting laboratories for generating the genetic sequence and metadata and sharing via the GISAID Initiative, on which this research is based (EPI_SET_260612ag GISAID identifier). We are very grateful for the field observations made by F. Lamo, E. Grøningssæter and H. Fålun Strøm. We also thank L.T. Engerdahl and I.A. Heffernan for their excellent work with diagnostics at NVI. We also sincerely thank K. Kovacs and C. Lydersen at NPI for communicating field observations to NVI.

The study was co-funded by the project OH4Surveillance, which is funded by the European Union (Grant Agreement No 101132473). Views and opinions expressed are those of the authors only and do not necessarily reflect those of the European Union or HaDEA. Neither the European Union nor the granting authority can be held responsible for them. Funding for this study was also provided by the Morris Animal Foundation (grant ID# D25ZO-430; KJB) and the Norwegian Veterinary Institute (SvalVilt, 12311).

Wild animals were captured, handled, and sampled by trained personnel using methods designed to minimize stress and risk of injury. Animals were released at the site of capture immediately after sampling. All procedures involving animals were conducted in accordance with national regulations and institutional guidelines (ARRIVE) for the care and use of animals. The study was approved by the Norwegian Animal Research Authority under permit number FOTS ID 31180.

## About the author

Dr. Madslien is a wildlife veterinarian, senior scientist and coordinator of wildlife health at NVI in Ås, Norway. His research has focused on terrestrial wildlife health for more than two decades.

