## Supplementary figures and images for "Highly Pathogenic Avian Influenza H5N5 in a Polar Bear and Atlantic Walrus, Svalbard, 2026, with Widespread Seroconversion in Polar Bears"

### Appendix Figure 1

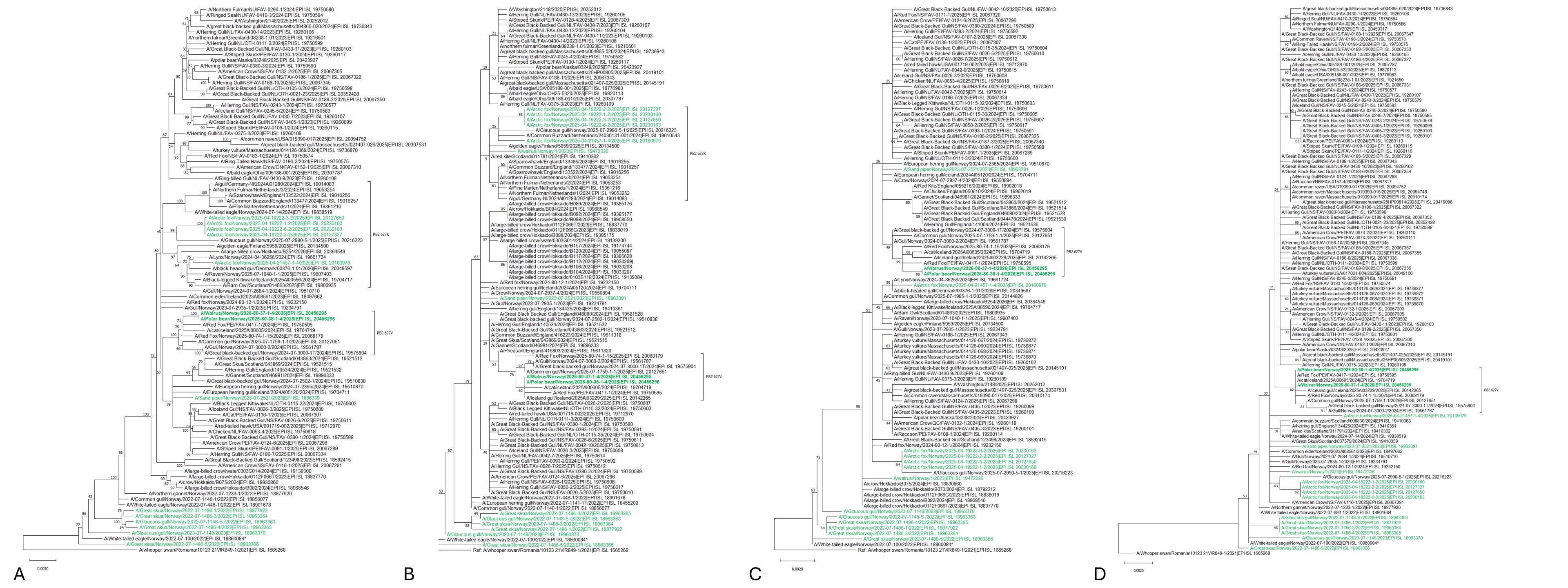
